# Aucubin promoted neuron functional recovery by suppressing inflammation and neuronal apoptosis in a spinal cord injury model

**DOI:** 10.1101/2022.02.01.478641

**Authors:** Shining Xiao, Nanshan Zhong, Quanming Yang, Anan Li, Weilai Tong, Yu Zhang, Geliang Yao, Shijiang Wang, Jiaming Liu, Zhili Liu

## Abstract

Spinal cord injury (SCI) can cause severe motor impairment. Post-SCI treatment has focused primarily on secondary injury, with neuroinflammation and neuronal apoptosis as the primary therapeutic targets. Aucubin (Au), a Chinese herbal medicine, exerts anti-inflammatory and neuroprotective effects. The therapeutic effects of Au in SCI have not been reported. We showed that Au can promote functional recovery after SCI. Recovery may occur through the toll-like receptor 4 (TLR4)/nuclear factor kappa B (NF-κB) pathway to promote M2/M1 polarization in microglia and inhibit mitochondrial dysfunction to reduce neuronal apoptosis. These biochemical changes result in reduced secondary injury and facilitate axon regeneration. Therefore, Au may be a promising post-SCI therapeutic medication.

## 1. Introduction

Spinal cord injury (SCI) is a terrible neurological condition that usually results in lifelong paraplegia and is associated with substantial physical and financial suffering [1–3]. However, there are no effective treatments for SCI. The pathological process of SCI can be divided into primary and secondary injuries [4, 5]. Primary SCI, which includes *in situ* structural damage, vascular rupture in the lesion area, and local bleeding around the lesion area, is caused by traumatic events (e.g., vehicle accidents or falling from a building) [6–8]. Secondary injuries are a group of disorders that develop following the primary injury, including severe inflammation, apoptosis, oxidative stress, ischemia, hypoxia, and axon damage [8]. Repair of spinal cord tissue is hampered by secondary damage. Studies have shown that effective treatment of secondary damage is critical to SCI recovery [9, 10], and inflammation and apoptosis are essential regulatory targets for prevention of secondary injury [11].

Inflammation is a critical effector of the healing process, but chronic inflammation exacerbates secondary complications following SCI [12]. Microglia are central nervous system macrophages that can express pro-inflammatory (M1) and anti-inflammatory (M2) phenotypes [13]. When suffering from SCI, microglia can be activated quickly. Pro-inflammatory mediators (e.g., cyclooxygenase (COX)-2, inducible nitric oxide synthase (iNOS), tumor necrosis factor (TNF)-α, Interleukin (IL)-1, and interleukin (IL)-6 etc.) are released by activated M1 phenotype microglia, resulting in an inflammatory response [14]. At the same time, SCI results in disruption of the blood-spinal-cord barrier. This disruption results in infiltration of peripheral immune cells (e.g., macrophages) at the injury site, which aggravates the inflammatory response [15]. After SCI, secondary injury increases and hampers the rehabilitation of linked functions due to inflammation inside and outside the central nervous system [16]. Studies have shown that increasing the M2/M1 polarization ratio in microglia/macrophages after SCI [17], or transplanting M2 phenotype microglia/macrophages to the injured spinal cord [18, 19], can promote functional recovery following SCI.

Neuronal apoptosis results in neurological dysfunction and is associated with poor prognosis [20, 21]. Reducing neuronal apoptosis can promote neurorehabilitation, neuroplasticity, and axonal regeneration, and has become a strategy for treatment of SCI [22]. Recent studies have shown that reducing the level of neuronal apoptosis can improve the recovery of motor function in rats that experienced SCI [23, 24]. Therefore, promoting M1 to M2 polarization of microglia/macrophages and inhibiting neuronal apoptosis may promote functional recovery following SCI.

Studies have shown that anti-neuroinflammatory medications and anti-apoptotic therapies have been shown to improve behavioral functions in a variety of animal models of SCI [5, 11]. However, few medications can enhance the M2/M1 polarization ratio in microglia/macrophages and prevent neuronal death. Therefore, anti-inflammatory and neuroprotective natural Chinese herbal medications have been extensively studied. Aucubin (Au) belongs to the iridoid glycoside family and is found in a variety of plant species [25]. A number of studies have shown that Au exerts potent anti-inflammatory [26], anti-oxidant [27], and anti-tumor effects [28], and protects liver [29] and brain tissue [25]. However, treatment of rats with Au following SCI has not been studied. In this study, we showed that Au promoted functional recovery of rats who subjected to SCI. Besides, we evaluated the mechanisms by which Au regulated microglia/macrophage polarization in SCI and exerted neuroprotective effects.

## 2. Methods

### 2.1 SCI model establishment and therapy

In animal experiments, all protocols are consistent with the Animal Care and Use Committee of the First Affiliated Hospital of Nanchang University. We purchased 36 adult female SD rats weighing approximately 220g from Tianqin Biotechnology (Changsha, China). Then the rats were randomly divided into three groups: sham operation (Sham group), SCI caused by extrusion (SCI group), SCI with Au (40 mg/kg) treatment (Au group). We anesthetized the rat with 1% (w/v) sodium pentobarbital (40 mg/kg) before removing the lamina at the T9 level to expose the spinal cord and modeling the spinal cord injury. After that, we used a vascular clip (15g force, Oscar) to compress the spinal cord tissue for 1 minute to create a SCI model. The rats in the Au group were injected intraperitoneally with Au (40 mg/kg) every day following the SCI operation, whereas the other groups received the same amount of normal saline. Furthermore, bladder squeezing was performed on the rats in the spinal cord injury groups twice daily to allow for urination.

### 2.2 Behavioral evaluation

Motor function was assessed using the Basso, Beattie, and Bresnahan (BBB) scale, and footprint analysis. For the BBB scale, on days 0, 1, 3, 7, 14, and 28 after surgery we placed the rats in an open environment and three independent observers who were unaware of the experimental procedure observed hind limb movement. The test took approximately 3 minutes, and the results were evaluated on a scale of 0 to 21 points (totally paralyzed hindlimb to normal mobility) [4]. For footprint analysis, we dyed the forelimbs and hind limbs of rats with black ink and red ink, respectively, and then had them walk on white paper, and evaluated the imprints [30].

### 2.3 Cell viability evaluation

The CCK8 kit (Beyotime, China) was used to measure the toxicity of Au (HPLC purity > 98%, Yaji Biotechnology, China) in BV2 cells. BV2 cells were plated at a density of 5000 cells per well in a 96-well plate. After a 24-hour incubation period with Au at concentrations of 0, 0.1, 1, 10, 100, 500, or 1000 μM, the BV2 cells were incubated for 2 hours with the CCK-8 kit before being measured the optical density (OD) at 450 nm.

We then evaluated the effects of Au on TBHP (Sigma)-induced neuronal apoptosis. PC12 cells were pretreated for 12 hours with Au (0, 25, 50, or 100 μM) and subsequently treated for 4 hours with TBHP (0 or 100 μM). The cells were analyzed using the CCK8 kit and OD was measured at 450 nm.

### 2.4 Cell culture

The BV2 microglia and PC12 cells were purchased from Cell Biology (Shanghai, China). The cells were resuspended in high-glucose DMEM medium (BV2 cells) and 1640 medium (PC12 cells), each containing 10% FBS (Gibco, Thermo). The cells were then cultured at 37°C in a humidified incubator containing 5% CO_2_. Every two days, the new medium was replenished.

### 2.5 Live-dead cell staining

A Calcein-AM/PI (Solarbio, China) double-staining experiment was utilized to assess the effect of Au on TBHP-induced PC12 cells. PC12 cells were planted in 6-well plates. After that, Au (0 or 100 μM) was pretreated for 12 hours before being incubated for 4 hours with TBHP (100 μM). Following that, PC12 cells were gently washed three times with 1×Assay Buffer to eliminate any leftover active esterase, and then incubated with Calcein-AM and PI combination for 15 minutes at 37°C.The photos are then captured using a fluorescence microscope (ZEISS).

### 2.6 Immunofluorescence staining

A round glass slide with an 18-mm diameter was placed on a 12-well plate, and BV2 cells were seeded on the round glass slide, *in vitro* study. The cells were treated with Au (0 or 100 μM) for 12 hours, then incubated with or without LPS (1 μg/mL, Sigma) for 24 hours. The cells were then fixed with 4% paraformaldehyde (PFA) for 15 minutes then permeabilized with 0.5% Triton X-100 for 30 minutes. The slide was blocked for 1 hour at 37 °C with 5% bovine serum albumin (BSA), then cultured overnight at 4 °C with the following primary antibodies: anti-Arg-1 (CST), anti-iNOS (CST), anti-NF-κB p65 (CST), anti-MAP2 (CST), and anti-Bcl-2 (CST). On the next day, the nuclei were stained with DAPI for 5 minutes before incubation with the appropriate secondary antibodies for 1 hour at 37 °C. The slides were then sealed with nail polish. A fluorescent microscope (ZEISS) was employes to visualize the cells.

In order to collect spinal cord tissue slices for immunofluorescence staining, we chose some rats on the 3rd and 28th days after SCI. The rats were anesthetized with chloral hydrate, and the left ventricle was perfused with 0.9% NaCl before being fixed with 4% PFA. The lesion’s spinal cord tissue is then removed, dehydrated, and embedded. Following that, a longitudinal paraffin slices with 5μm thickness were prepared. Following antigen retrieval, paraffin slices were blocked for 1 hour at 37°C with 5% BSA before being treated overnight at 4°C with various primary antibodies as follows: anti-GFAP (Abcam), anti-CD68 (Abcam), anti-Arg-1 (CST), anti-NeuN (Abcam), anti-Cleaved Caspase 3 (CST), anti-MAP2 (CST) and anti-GAP43(CST). The following methods are identical to those used *in vitro*.

### 2.7 TUNEL staining

Cell apoptosis can be detected using TUNEL (Keygen Biotech, China) staining. We permeabilized paraffin sections with 0.5% Triton X-100 for 10 minutes at room temperature following antigen retrieval. The paraffin sections were then incubated with a mixture of labeling solutions (1X Labeling Solution and TdT), then with DAPI for nuclear staining. The cells were visualized using a fluorescence microscope (ZEISS).

### 2.8 Western blot

Total protein is isolated from cells and spinal cord tissue. The extract was quantified with BCA reagent and equilibrated to the same volume. The same amount of protein was loaded into the combs of different SDS-polyacrylamide gels before electrophoresis. Then transfer the protein bands to a polyvinylidene fluoride (PVDF) membrane, and block with 5% skimmed milk powder for 2 hours and then incubate overnight at 4°C with appropriate primary antibodies as follows: anti-TLR4 (CST), anti-MyD88 (Proteintech), anti-IκBα (CST), anti-p-IκBα (CST), anti-p65 (CST), anti-p-p65 (CST), anti-GAP43 (CST), anti-MAP2 (CST), anti-Arg-1(CST), anti-iNOS (Proteintech), anti-COX-2 (Abcam), anti-NeuN (Abcam), anti-Cleaved Caspase 3 (CST), anti-Bcl-2 (Proteintech), anti-BAX (Proteintech), anti-β-Actin (Proteintech) and anti-GAPDH (Abcam). After the next day, the bands were treated with the matching secondary antibodies for 2 hours. Subsequently, the band was detected by the imaging system (Tanon), and the ImageJ software was used for quantitative analysis.

### 2.9 Histological staining

Paraffin slices (5 μm thickness) were dewaxed, then stained with hematoxylin and eosin (H&E) as described previously [31]. All images were captured using an optical microscope (ZEISS).

### 2.10. Statistical analysis

The mean ±standard deviation was used to express all results. The data from each group were analyzed using GraphPad Prism 7.0. Statistical significance was determined by one-way analysis of variance (ANOVA) followed by Tukey’s multiple comparisons. P <0.05 indicates statistical significance.

## 3. Results

### 3.1 Au promoted recovery of motor function in rats following SCI

Behavior (footprint test and BBB locomotion score) and pathology were used to assess functional recovery following SCI. Hind limb trajectories were strikingly different among the groups in the footprint test. After 28 days, compared to the clear footprints of the Sham rats, the hind limbs of the SCI rats showed two obvious drag marks (black ink), while the SCI rats that received Au therapy showed some coordinated movement, as evidenced by several black footprints (red arrows; Fig. 1D). Moreover, the BBB score showed that the rats that were subjected to SCI lost motor function on the first day after surgery, and function of the hind limbs returned gradually. Interestingly, the Au therapy group’s movement of the hind limbs was clearly superior to the SCI group’s (Fig. 1E). Staining with H&E at 28 days showed that the cystic cavity was much smaller in the Au treatment group than that in the SCI group (Fig. 1F and G). These results indicated that Au may improve pathology and motor function in rats that are subjected to SCI.

**Fig. 1.**
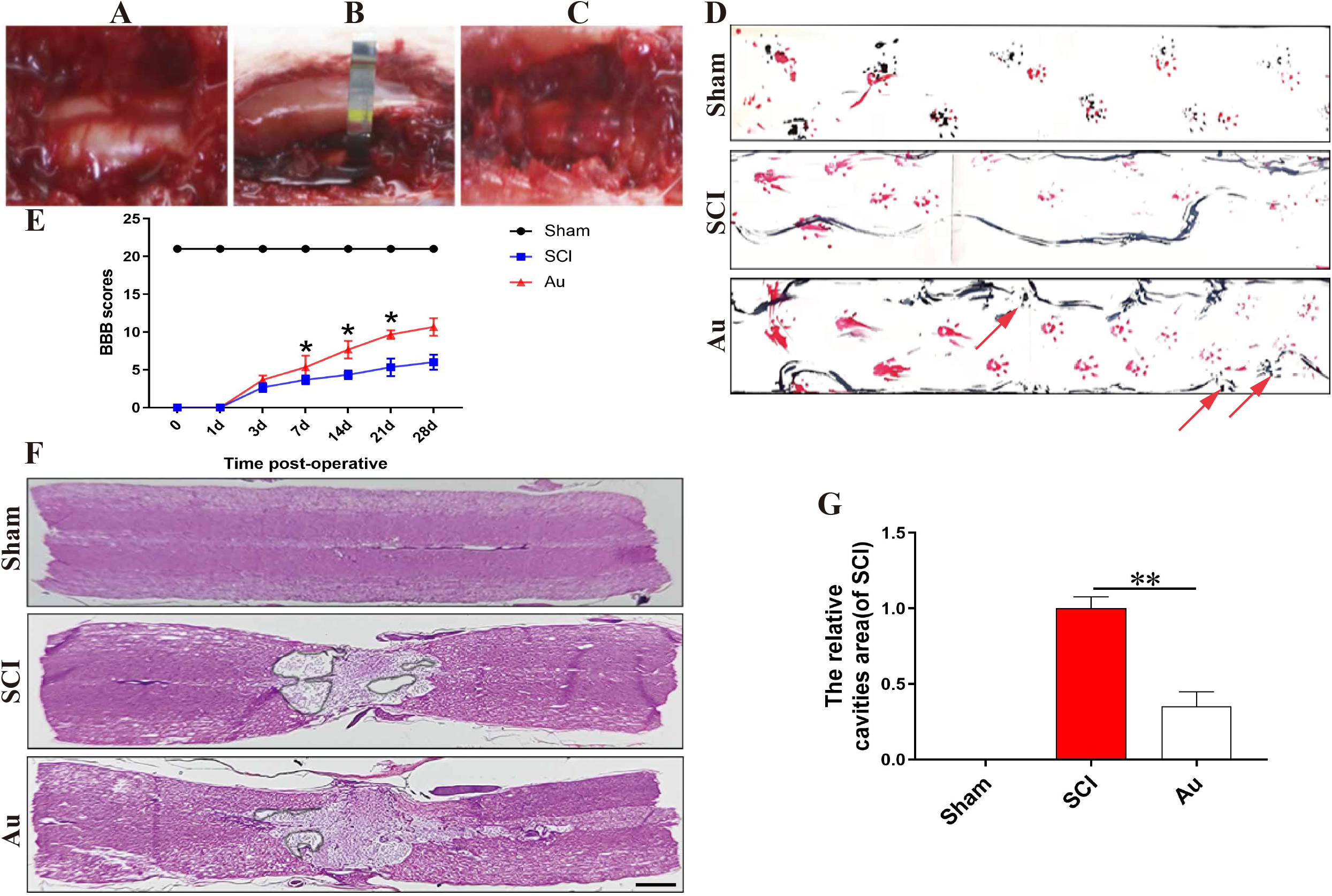
Au improved motor function after SCI and reduced cavity area. (A) The spinal cord of a rat that has been exposed. (B) A vascular clip is used to compress the exposed spinal cord. (C) The site of the crushing. (D, E) The behavior assay of footprint assay and BBB scores at 28 days postinjury (red arrow pointed the posterior limb prints). (F, G) H&E staining was used to calculate the extent of the lesion cavity in each group at 28 days postinjury (scale bar =1000μm). N ≥ 3 per group for separate experiments. **P and ***P indicate Au group versus SCI group. *P < 0.05 and **P < 0.01.

### 3.2 Au inhibited the inflammatory response after SCI by promoting M2/M1 polarization in microglia/macrophages

To explore the effects of Au on microglia/macrophage polarization after SCI, we selected rats on the third day after surgery for evaluation. Immunofluorescence staining showed that CD68 microglia/macrophages (M1 phenotype) in the SCI group were concentrated in the injured area and in the surrounding normal (green), and Au treatment visibly reduced the number of activated microglia/macrophages in the lesion area and in the surrounding tissue (Fig. 2A and C). Additionally, the protein levels of CD68, iNOS, and COX-2 after Au treatment were significantly lower than those in the SCI group (Fig. 2D-H), which indicated that Au inhibited polarization to the M1 phenotype and reduced the inflammatory response following SCI. To further evaluate M2 polarization after SCI, we used Arg-1 (a M2 microglia/macrophage phenotype marker) immunofluorescence staining to evaluate the distribution of positive cells in the injured area. The results showed that compared with the SCI group, the Sham group hardly saw M2 phenotype microglia/macrophage, while the Au treatment group had more related M2-positive cells (Fig. 2B and C). Furthermore, high Arg-1 protein expression in the Au group also showed that Au treatment promoted M2 polarization (Fig. 2D and E). These results may have been due to the presence of a small population of M2 polarized microglia, even following SCI [11, 32]. Therefore, the number of M2 microglia/macrophages in the SCI group was higher than that in the Sham group. Furthermore, the infiltration of immune cells into the injured area increased, and Au treatment significantly promoted M2 polarization, which resulted in a greater number of M2 polarized microglia/macrophages in the Au treatment group. These results indicated that Au promoted M2/M1 polarization after SCI, which resulted in decreased inflammation.

**Fig 2.**
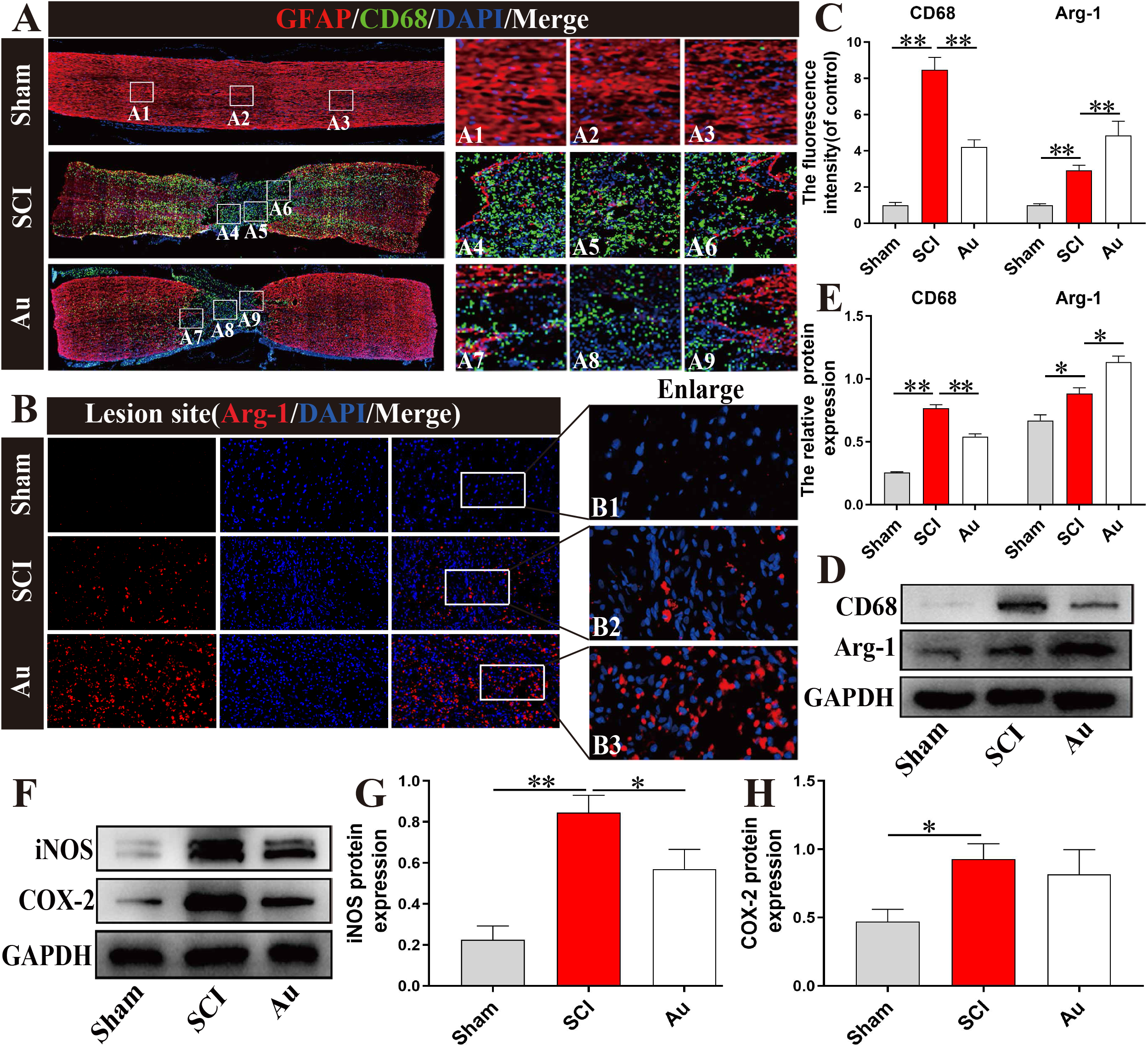
Au treatment suppresses the inflammatory response after SCI. (A, C) CD68 (green)/GFAP (red) double fluorescence labeling and quantitative examination of spinal cord tissue sections in each group at 3 days postinjury (scale bar=1000μm). A1, A4 and A7 are the enlarged views of the solid white box area from the rostral area. A2, A5 and A8 are the enlarged views of the solid white box area from the lesion area. A3, A6 and A9 are the enlarged views of the solid white box area from the caudal area. (B, C) Representative images containing Arg-1 staining and quantification analysis in each group at 3 days after SCI (scale bar=100μm). B1, B2 and B3 are the enlarged figures of the solid white box area. (D, E) CD68 and Arg-1 proteins were detected and quantified in each group at 3 days postinjury. (F, G, H) iNOS and COX-2 proteins were detected and quantified in each group at 3 days postinjury. N ≥ 3 per group for separate experiments. *P < 0.05 and **P < 0.01.

### 3.3 Au increased the M2/M1 polarization ratio in microglia

We first evaluated Au toxicity in BV2 cells. As shown in Figure 3A, treatment of BV2 cells with different concentrations of Au (0, 0.1, 1, 10, 100, 500, and 1000 μM) for 24 hours resulted in no apparent toxicity below 100 μM. Viability of BV2 cells was significantly lower following treatment with 500 μM or 1000 μM Au. Therefore, we treated cells with 100 μM or less.

**Fig. 3.**
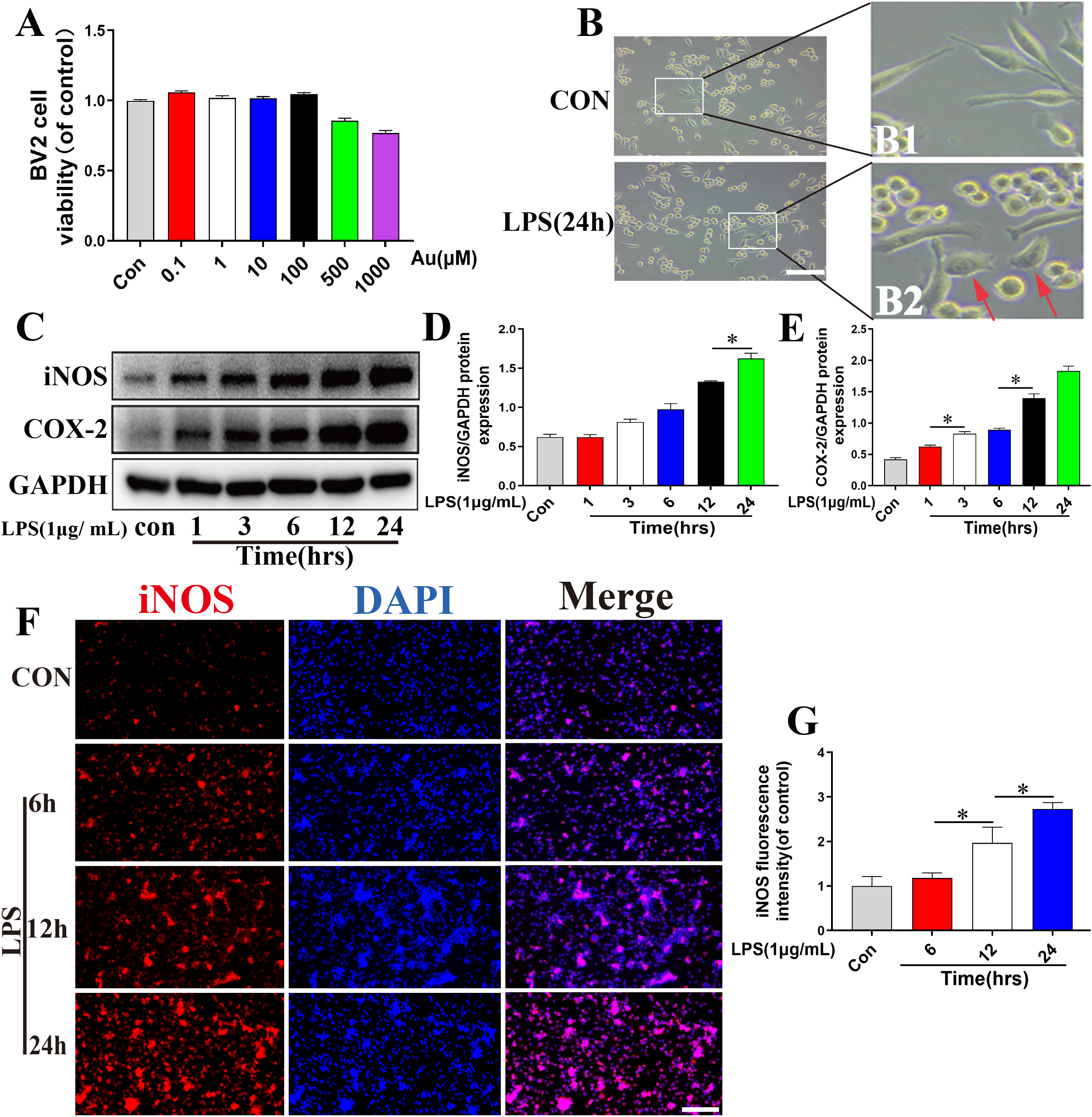
M1 microglia polarization was triggered by LPS in a time-dependent manner. (A) The CCK-8 test was applied to assess the viability of BV2 cells treated with Au at various doses for 24 hours. (B) Morphological change of BV2 cells (scale bar=100μm). B1 and B2 are the enlarged figures of the solid white box area (red arrow pointed the amoebic change). (C, D, E) iNOS and COX-2 proteins were detected and quantified in each group of microglia. (F, G) iNOS level was measured by immunofluorescence labeling and quantification examination in each group of microglia (scale bar=200μm). N ≥ 3 per group for separate experiments. *P < 0.05.

Next, we studied the effect of LPS (1 μg/mL) induction on release of pro-inflammatory mediators *in vitro*. An amoebic alteration (Fig. 3B) in cell shape (red arrow indicated) was observed after LPS stimulated BV2 cells for 24 hours, which showed that BV2 cells were activated to the M1 phenotype. Western blot results showed that the expression levels of pro-inflammatory mediators (e.g., iNOS and COX-2) in BV2 cells after LPS treatment increased in a time-dependent manner within 24 hours (Fig. 3C-E). Furthermore, immunofluorescence staining of iNOS (Fig. 3F and G) agreed with the western blot results. As iNOS and COX-2 are M1 microglial markers, these results indicated that stimulation of BV2 cells with LPS promoted M1 polarization with a peak at 24 hours.

To investigate the influence of Au on microglia polarization, BV2 cells were pretreated with Au for 12 hours, then stimulated with LPS (1 μg/mL) for 24 hours. Figure 4A-C shows that pretreatment with Au (25, 50, and 100 μM) reduced the LPS-induced increases in iNOS and COX-2 levels, which agreed with the results of iNOS immunofluorescence analysis (Fig. 4D and E). The results indicated that Au could prevent M1 polarization of microglia. Western blot and immunofluorescence analysis showed that Au pretreatment increased the expression of M2 microglia-related markers (Arg-1) (Fig. 4G-I). In summary, these results indicated that Au inhibited M1 polarization of microglia and promoted the level of polarization of M2 phenotype microglia in a dose-dependent manner.

**Fig. 4.**
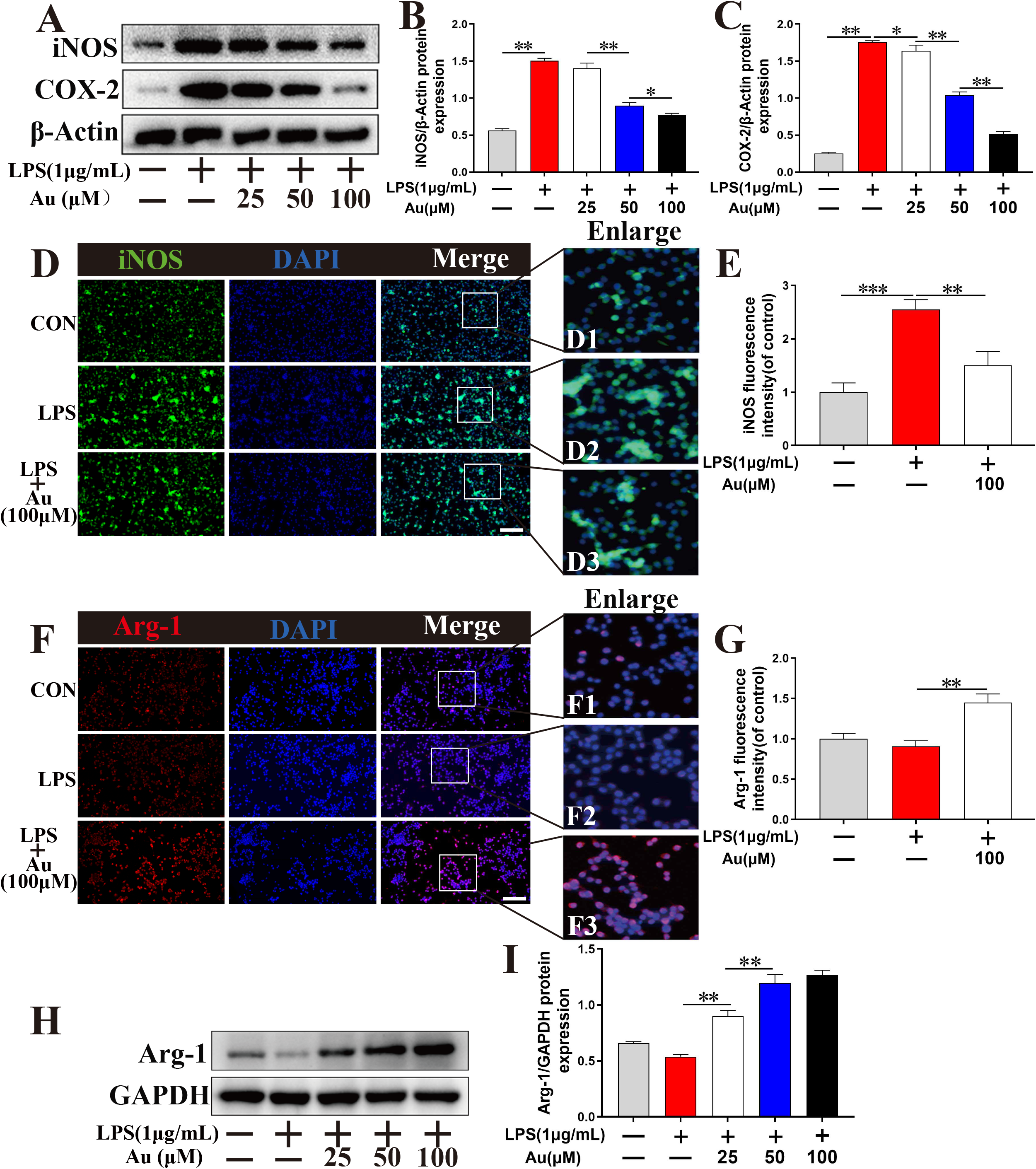
Au promotes M2/M1 cell polarization ratio in microglia. (A, B, C) iNOS and COX-2 proteins were detected and quantified in microglia. (D, E) Immunofluorescence staining and quantitative data for iNOS in each group of microglia (scale bar=200μm). D1, D2 and D3 are the enlarged figures of the solid white box area. (F, G) Immunofluorescence staining and quantification examination revealed the expression levels of Arg-1 in each group of microglia (scale bar=200μm). F1, F2 and F3 are the enlarged figures of the solid white box area. (H, I) Arg-1 protein was detected and quantified in each group of BV2 cells. N ≥ 3 per group for separate experiments. *P < 0.05, **P < 0.01 and ***P < 0.001.

### 3.4 Au inhibited release of LPS-induced pro-inflammatory mediators by regulating the TLR4/NF-κB pathway

The toll-like receptor 4 (TLR4)-myeloid differentiation protein-88 (MyD88) signaling pathway has been closely related to inflammation in numerous studies [33, 34]. To determine whether Au suppressed LPS-induced microglial activation via the TLR4-mediated MyD88 pathway, we pretreated BV2 microglia with Au (0, 25, 50, and 100 μM) for 12 hours before stimulating them with LPS (1 μg/mL) for 24 hours. Treatment with LPS resulted in increased protein expression levels of TLR4 and MyD88 in BV2 microglia, as determined using western blot. Treatment with Au reversed LPS-induced expression of TLR4 and MyD88 in a concentration-dependent manner (Fig. 5A-C). These findings were consistent with the anti-inflammatory effects of Au (Fig. 4A-C). These results indicated that Au reduced inflammation through inhibition of the TLR4-MyD88 pathway.

**Fig. 5.**
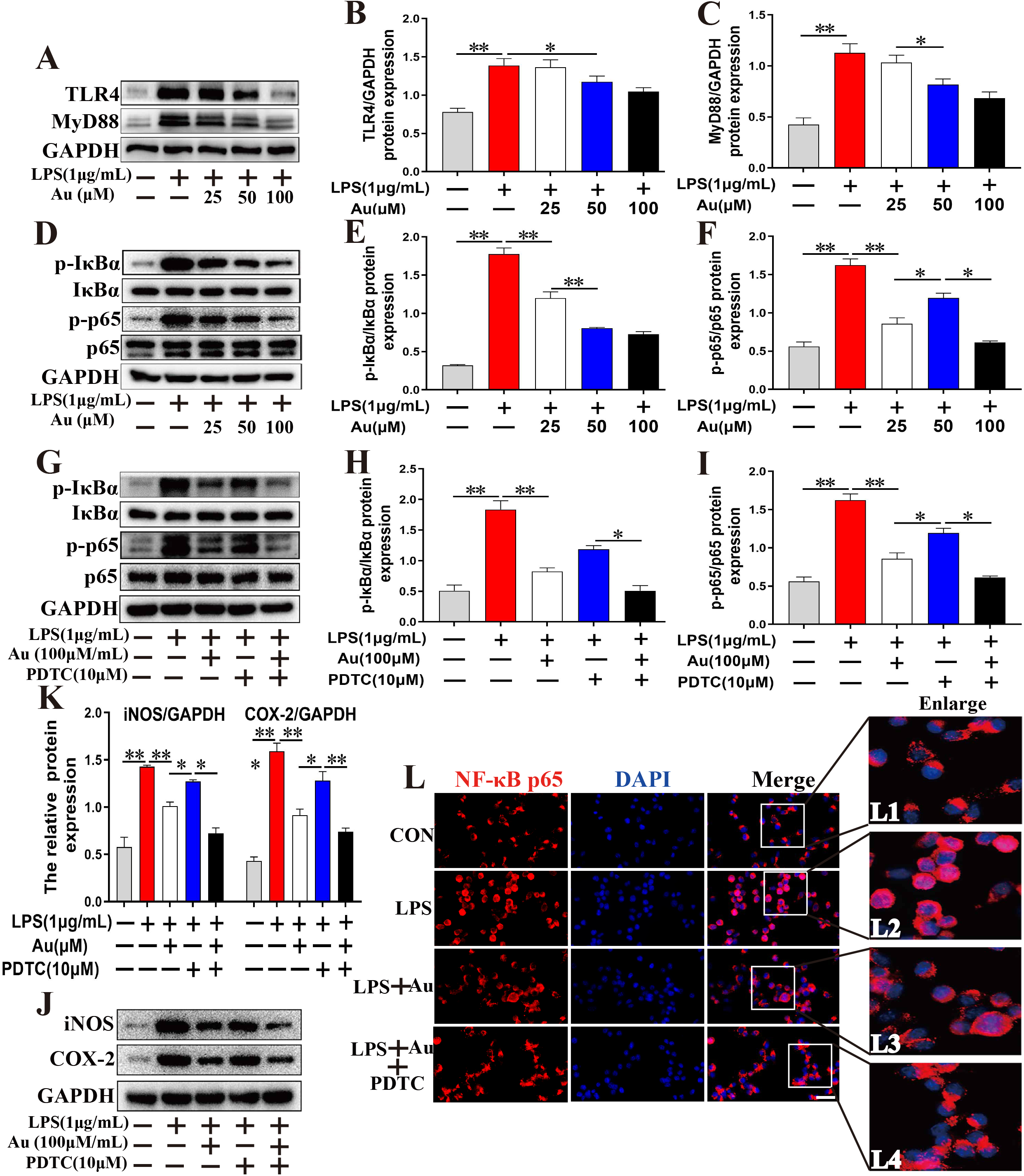
Au treatment suppresses inflammatory response via TLR4/NF-κB pathways *in vitro*. (A, B, C) TLR4 and MyD88 proteins were detected and quantified in each group of microglia. (D, E, F, G, H, I) Protein expressions IκBα, p-IκBα, p65 and p-p65 in each group of microglia. (J, K) iNOS and COX-2 proteins were detected and quantified in each group of microglia. (L) The translocation of the NF-κB p65 was determined by immunofluorescence staining in each group of microglia. (scale bar=50μm). L1, L2 and L3 are the enlarged figures of the solid white box area. N ≥ 3 per group for separate experiments. *P < 0.05 and **P < 0.01.

Furthermore, studies have shown that the pro-inflammatory response mediated by the TLR4-MyD88 signaling pathway is linked to a number of other signaling pathways, including nuclear factor kappa B (NF-κB) pathway [35, 36]. We evaluated whether Au inhibited the release of pro-inflammatory mediators in microglia via the NF-κB pathway. Western blot showed that LPS greatly increased the phosphorylation levels of inhibitor kappa Bα (IκBα) and p65 in microglia, and Au dramatically suppressed this effect (Fig. 5D-F). We used PDTC, an NF-κB pathway inhibitor, to further investigate the role of NF-κB signaling in Au-mediated reductions in release of pro-inflammatory mediators. BV2 microglia were pretreated for 2 hours with or without PDTC (10 μM), treated for 12 hours with Au (25, 50, 100 μM), then cultured in LPS-containing medium for 24 hours. As shown in Figure 5G-I, treatment with PDTC and Au inhibited phosphorylation of IκBα and p65 more than that observed with PDTC or Au alone. We also used Western blot to observe if Au affects NF-κB signaling to alter the LPS-induced pro-inflammatory response. When compared to the effects of PDTC or Au alone, co-incubation with PDTC and Au visibly reduced iNOS and COX-2 protein expression levels (Fig. 5J and K). These findings were further validated by immunofluorescence staining, which showed that the combination of PDTC and Au effectively prevented LPS-induced nuclear translocation of NF-κB p65 to the nucleus (Fig. 5L). These results demonstrated that the anti-inflammatory effects of Au occurred through modulation of the NF-κB pathway.

### 3.5 Au reduced neuronal apoptosis *in vivo* and *in vitro*

Studies have shown that reducing neuronal apoptosis promoted functional recovery after SCI [37–39]. We found that administration of Au reduced cell apoptosis in SCI, as determined using TUNEL staining (Fig. 6A and B). Furthermore, western blot analysis showed that Au treatment partially restored NeuN (neuronal marker) protein expression (Fig. 6C and D). These results indicated that Au treatment inhibited SCI-induced neuronal apoptosis. We further evaluated the effect of Au on neuronal apoptosis through evaluation of expression of apoptotic proteins. Compared with the Sham group, the SCI group had distinctly increased expression of mitochondrial pro-apoptotic proteins (Bax and cleaved caspase-3), and decreased expression of mitochondrial anti-apoptotic proteins (Bcl-2). Treatment with Au partially reversed these changes in protein expression (Fig. 6E and F). Staining for cleaved caspase-3 produced results that agreed with those obtained by western blot (Fig. 6G and H). These results indicated that administration of Au could reduce neuronal apoptosis by improving mitochondrial dysfunction after SCI.

**Fig. 6.**
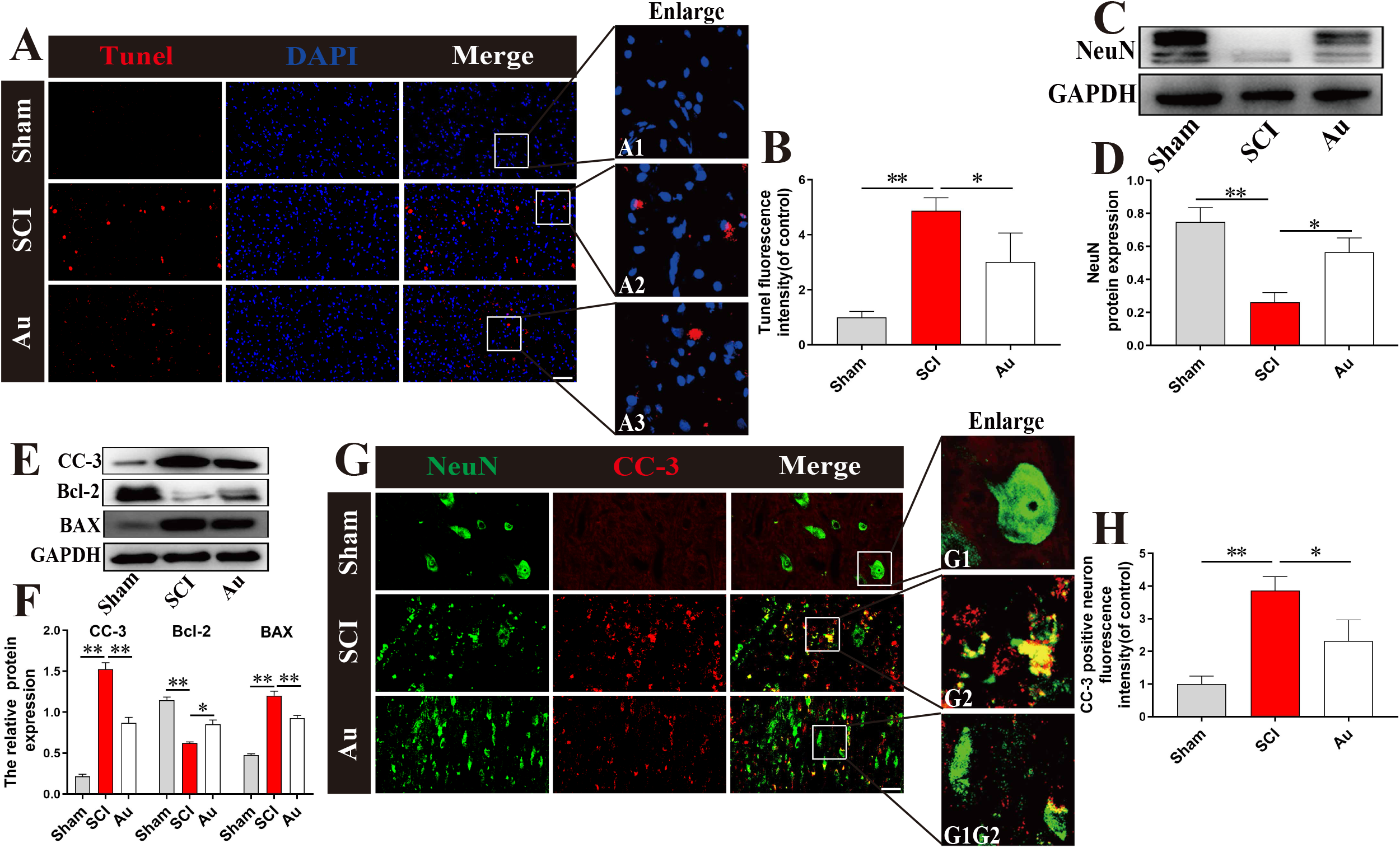
Pretreatment with Au prevents apoptosis in spinal neurons after SCI. (A, B) Tunel staining and quantitative analysis were used to detect neuronal apoptosis in each group at 3 days postinjury (scale bar=100μm). A1, A2 and A3 are the enlarged figures of the solid white box area. (C, D) NeuN protein level were detected and quantified in each group at 3 days postinjury. (E, F) Apoptosis-related levels (CC-3, Bcl-2 and BAX) were detected and quantified in each group at 3 days postinjury. (G, H) Representative images and quantification analysis of CC-3 (red) and NeuN (green) fluorescence staining in each group at 3 days postinjury (scale bar=50μm). G1, G2 and G3 are the enlarged figures of the solid white box area. N ≥ 3 per group for separate experiments. *P < 0.05 and **P < 0.01.

To further evaluate the effects of Au on apoptosis, PC12 cells were treated with TBHP to induce oxidative stress and to trigger neuronal apoptosis. Cell viability was determined using the CCK8 kit, and the results showed that Au treatment significantly reversed TBHP-induced neuronal apoptosis (Fig. 7B). We then measured the levels of the mitochondrial anti-apoptosis-related protein Bcl-2. The results showed that Bcl-2 expression was evidently lower in the TBHP group than that in the CON group. However, Bcl-2 level was partially recovered in the Au treatment group (Fig. 7C and D). Furthermore, of the results of staining for Bcl-2 agreed with those obtained using western blot (Fig. 7E and F). Double-staining with calcein-AM and PI also showed that Au inhibited TBHP-induced apoptosis in PC12 cells (Fig. 7G and H). Microscopic analysis showed that PC12 cells in the TBHP group underwent early apoptosis (permeability was clearly increased, and the cell bodies and axons had contracted), and cell shape was partially restored following Au treatment (Fig. 7I). These results indicated that Au could reverse neuronal apoptosis and enhance neuronal survival by improving mitochondrial dysfunction *in vivo* and in *vitro*.

**Fig. 7.**
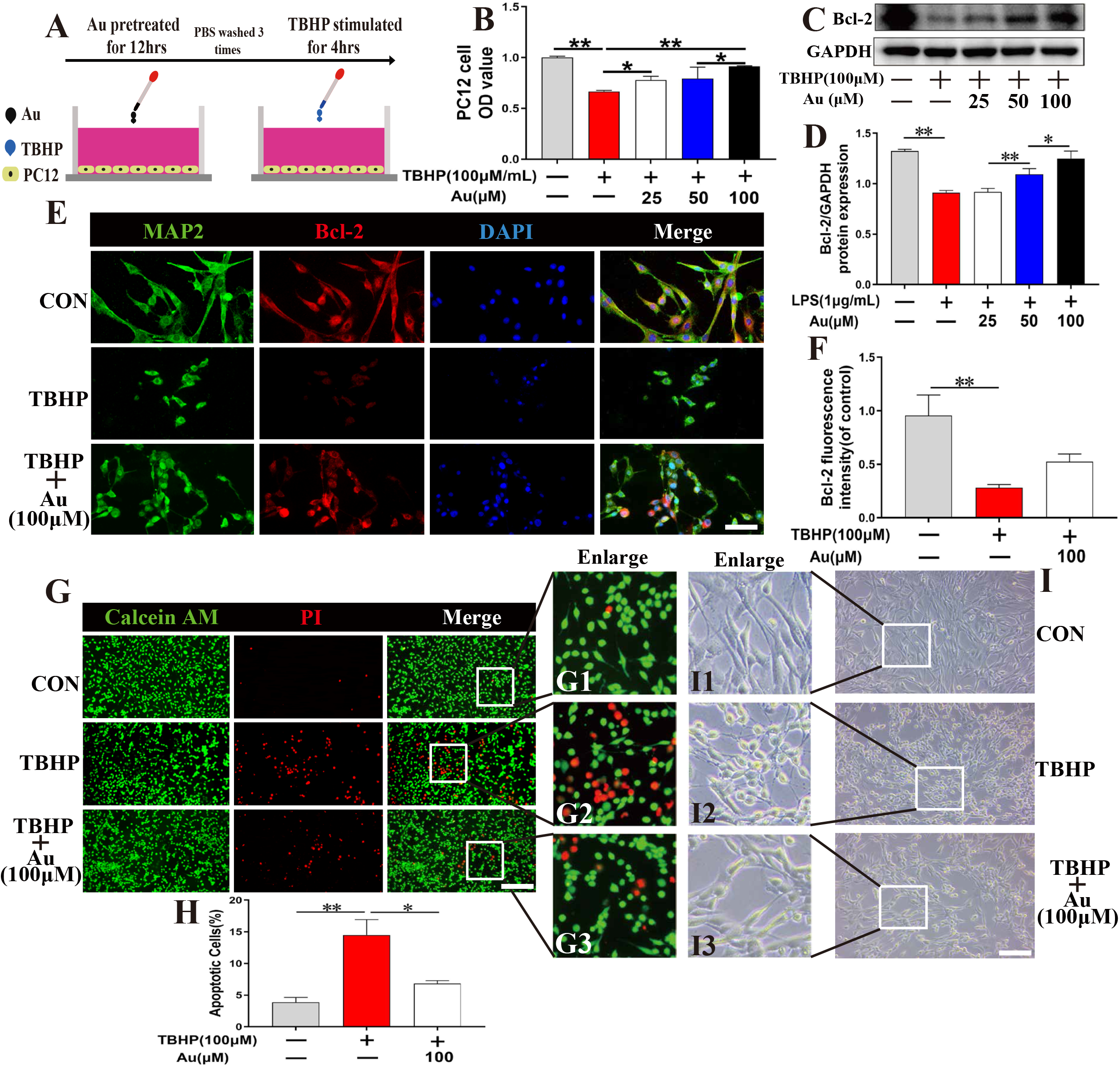
Au reduces neuronal apoptosis induced by TBHP. (A) Treatments for PC12 cells are depicted in this diagram. PC12 cells were pretreated for 12 hours with Au, then washed three times before being stimulated with TBHP for 4 hours. (B) The effect of Au pretreatment on the viability of PC12 cells was evaluated by CCK-8 test in each group. (C, D) Representative western blotting images and quantification showing Bcl-2 protein levels in each group of PC12 cells. (E, F) MAP2 and Bcl-2 immunolabeling and quantitative analysis in each group of neurons (scale bar=50μm). (G-H) Double labeling with Calcein AM/PI and quantitative examination of neuronal apoptosis in each group of PC12 cells (scale bar=200μm). G1, G2 and G3 are the enlarged figures of the solid white box area. (I) Morphological change of PC12 cells in each group (scale bar=200μm). I1, I2 and I3 are the enlarged figures of the solid white box area. N ≥ 3 per group for separate experiments. *P < 0.05, **P < 0.01 and ***P < 0.001.

### 3.6 Au promoted axon regeneration after SCI in rats

We measured the expression of an axon microtubule component, MAP2, in spinal cord tissue using immunofluorescence staining. The rostral and caudal axons in the SCI group showed sparse distribution and low-intensity staining. In contrast, Au treatment resulted in a more regular distribution of MAP2-positive axons, with increased density and staining intensity, and evidence of axonal growth toward the injured area (Fig. 8B and C). These results showed that Au promoted axon regeneration and extension following SCI. Additionally, we evaluated extension of nerve filaments in the injured spinal cord by GAP43 staining. Compared with the SCI group, the Au group had significantly higher expression of GAP43 nerve fibers at the rostral and caudal regions, and these nerve fibers appeared as more punctate or tubular (Fig. 8D and E). Furthermore, the expression levels of MAP2 and GAP43 showed that Au treatment reversed axonal damage induced by SCI (Fig. 7F-H). These results demonstrated that Au can promote axon regeneration in damaged spinal cord *in vivo*.

**Fig. 8.**
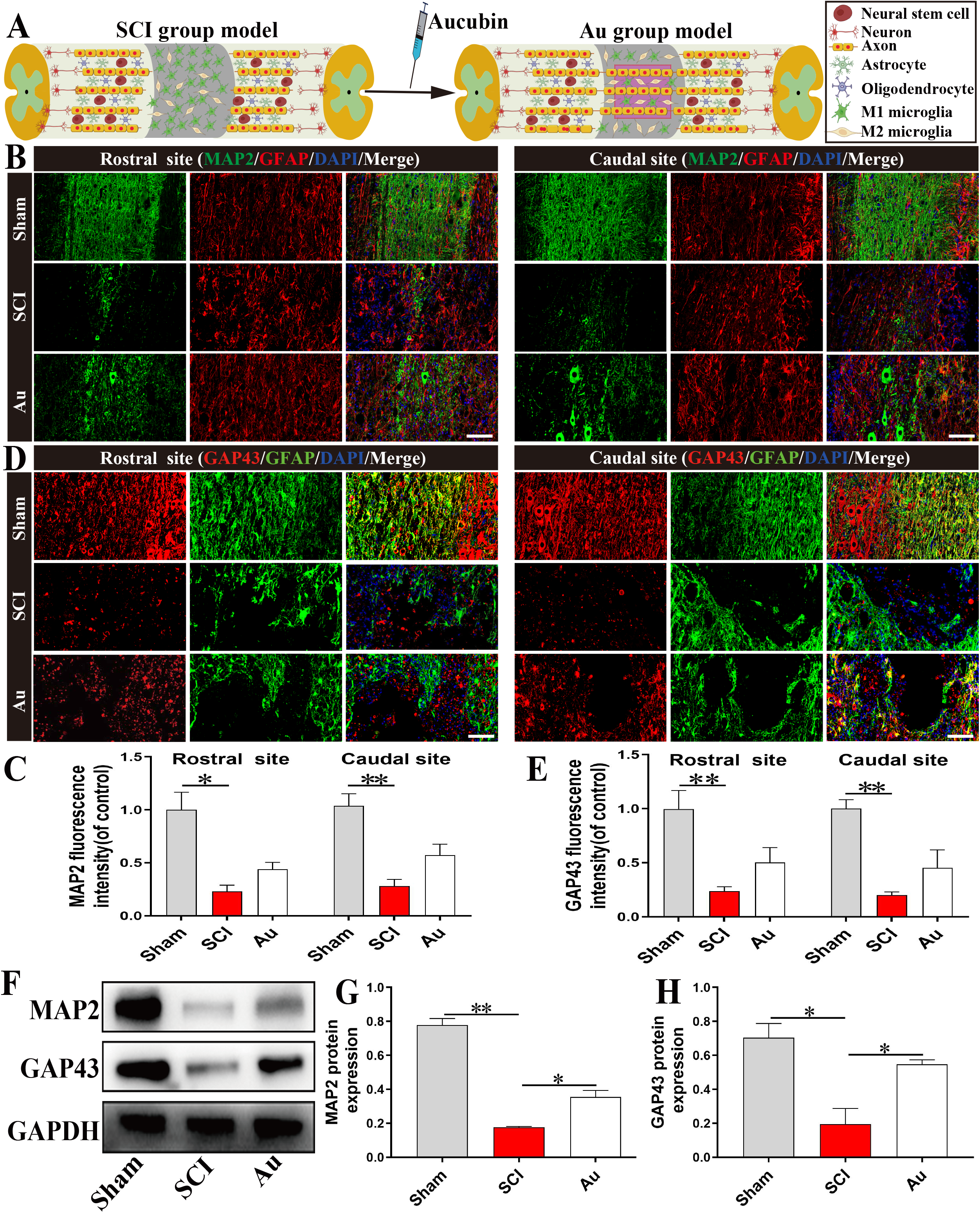
After SCI, Au improves neuronal survival and axonal regeneration. (A) Schematic diagram showing that Au improves axon regeneration. (B, C) At 28 days after SCI, double immunofluorescence pictures and quantification examination revealed the expression levels of MAP2 (green) and GFAP (red) in each group (scale bar=100μm). (D, E) Representative images containing neurofilament (GAP43) staining and quantitative analysis in each group at 28 days postinjury (scale bar = 100μm). (F, H) MAP2 and GAP4 proteins were detected and quantified in each group at 28 days postinjury. N ≥ 3 per group for separate experiments. *P < 0.05 and **P < 0.01.

**Fig 9.**
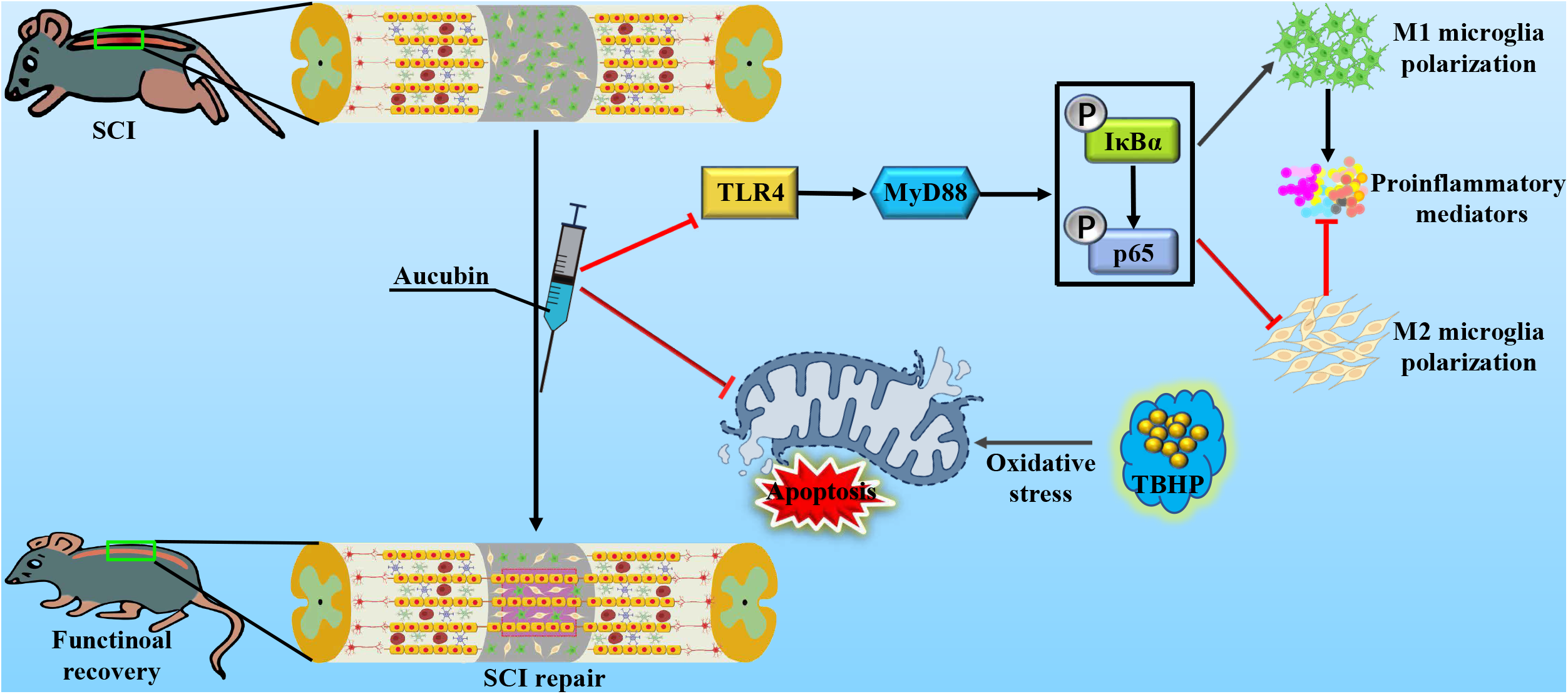
Au’s therapeutic effects on SCI are depicted in this diagram. Au can increase the M2/M1 cell ratio by inhibiting the TLR4/NF-κB pathway, suppress neuronal apoptosis via inhibit mitochondrial dysfunction, and ultimately promote the functional recovery of rats with SCI.

## 4. Discussion

SCI is a severely debilitating condition that results in significant loss of sensory and motor function [40–42]. Extensive research has focused on identification of medications that can promote functional and pathological recovery following spinal cord injury [43–45]. We studied a Chinese herbal medicine called Au, which has a strong therapeutic effect on SCI, and explained the mechanism by which Au promoted recovery of motor function in rats with SCI. The average hindlimb score of the SCI group was about 6 points, while the average hindlimb score in the Au group was about 12 points (Fig. 1E). These results were consistent with the general trend of the footprint analysis results (Fig. 1D). This finding demonstrated that Au promoted rehabilitation of hindlimb motor function following SCI. Furthermore, H&E staining showed that Au reduced the area of cystic cavities compared to that in the SCI group (Fig. 1F and G), showing that Au aided functional recovery following SCI.

Secondary injury, which typically manifests as severe inflammation and neuronal apoptosis [11, 46], is the principal pathogenic process that aggravates tissue damage and hampers recovery of motor function following SCI [4, 47]. Because secondary injury is a critical therapy window, several studies have focused on reducing inflammation and neuronal death following SCI to aid functional recovery [48, 49]. Microglia become over-activated if the central nervous system is injured, resulting in the release of a large number of inflammatory mediators [50, 51]. Generation of these mediators is uncontrolled, and is accompanied by the release of neurotoxic substances such as free radicals and acute-phase proteins, and infiltration of inflammatory cells [43, 52]. Inflammatory mediators and chemokines worsen secondary damage, resulting in a negative impact on recovery following SCI [12, 53]. A large number of studies have shown that reducing central nervous system inflammation and peripheral inflammatory cell infiltration (e.g., macrophage) can significantly reduce secondary damage and promote functional recovery following SCI [16, 43]. After SCI, microglia/macrophages not only polarize into M1 cells causing inflammation, but they may also partially polarize into M2 cells, which have anti-inflammatory properties [5, 54]. The polarized orientation of microglia/macrophages after SCI is strongly skewed toward the M1 phenotype [16]. Studies have shown that modulating the polarization of microglia/macrophages between M1 and M2 following SCI can significantly impact the inflammatory response [5, 11]. Therefore, promoting M2 polarization of microglia/macrophages in the injured spinal cord may significantly improve SCI recovery. After SCI, the number of M1 phenotype microglia/macrophages increased at the injury site and permeated the surrounding area. Surprisingly, the number of M1 cells in the Au group was significantly reduced and mostly restricted to the lesion area (Fig. 2A and C), while the protein levels of the relevant M1 cell markers (CD68, iNOS, and COX-2) were also clearly reduced (Fig. 2D-H). More notably, the Au group had much greater positive numbers and protein levels of M2 microglia/macrophage than the SCI group (Fig. 2B-E). In addition, we used LPS to activate BV2 cells *in vitro* to mimic neuroinflammation. The results showed that LPS treatment polarized BV2 cells to the M1 phenotype and enhanced release of pro-inflammatory mediators, with a peak at 24 hours (Fig. 3). Pretreatment with Au reduced LPS-induced M1 polarization of BV2 cells and promoted M2 polarization (Fig. 4). These results were consistent with the effects of Au treatment on microglia/macrophages *in vivo*. We showed that Au treatment increased the M2/M1 ratio for microglia/macrophages and reduced inflammation *in vivo* and *in vitro*.

Toll-like receptor 4, a member of the toll-like receptor family, plays an important role in cellular immunity [55]. Studies have shown that TLR4 is abundantly expressed in microglia and is associated with pathogenesis of neuroinflammation [56]. Toll-like receptor 4 can bind to various signaling proteins, such as MyD88, to activate downstream signals, resulting in immune system activation and increased expression of inflammatory genes [57]. We showed that LPS increased TLR4 and MyD88 expression in microglia. In contrast, Au clearly reversed the LPS-induced increase in TLR4 and MyD88 levels (Fig. 5A-C). These data were consistent with results showing that Au inhibited release of pro-inflammatory mediators from microglia *in vitro* (Fig. 4A-C). These results suggest that Au blocked the TLR4-MyD88 signaling pathway, thereby inhibiting the neuroinflammation in LPS-activated microglia. The TLR4-mediated pro-inflammatory response is also linked to a number of signaling pathways, including the NF-κB pathway [36], which is a key component in the inflammatory response [58]. Studies have shown that IκBα is degraded once the NF-κB pathway is activated, which increases the phosphorylation of p65, allowing p65 to translocate from the cytoplasm to the nucleus, resulting in inflammation [59]. We found that Au not only inhibits the expression of p-IκBα, but also inhibits p65 phosphorylation and regulates its nuclear translocation (Fig. 5D, E, F, and L), indicating that Au inhibits the NF-κB pathway. To investigate whether Au suppressed LPS-induced microglial activation via NF-κB signaling, we treated microglia with PDTC, an NF-κB pathway inhibitor. The results showed that Au or PDTC therapy lowered the protein levels of p-IκBα, p-p65, iNOS, and COX2. Furthermore, co-treatment with Au and PDTC significantly reduced the expression of p-IκBα, p-p65, and inflammatory markers (e.g., iNOS and COX2) (Fig. 5G-K). Fluorescence results showed that co-incubation with Au and PDTC resulted in significantly greater translocation of p65 to the nucleus compared with that observed in response to Au or PDTC incubation alone (Fig. 5L). Our findings showed that Au exerted anti-inflammatory properties by inactivating the NF-κB pathway, which was mediated by TLR4-MyD88.

Neuronal apoptosis is an important component of secondary injury [37]. Inhibition of neuronal apoptosis is thought to aid in neural rehabilitation and axon regeneration [39, 60]. Therefore, development of strategies to reduce neuronal apoptosis in SCI has received greater interest. We used TUNEL staining to measure the level of cell apoptosis after acute SCI, and the results showed that Au therapy dramatically reduced cell apoptosis (Fig. 6A and B). Furthermore, the protein level of NeuN (a neuron marker) was higher in the Au group than that in the SCI group (Fig. 6C and D). These results suggested that administration of Au after SCI inhibited neuronal apoptosis. Additionally, we found that, compared to the SCI group, the levels of the mitochondrial apoptosis-related proteins (BAX and Cleaved Caspase 3) rose in the SCI group, while the expression levels of the mitochondrial anti-apoptotic protein Bcl-2 decreased; however, this result was reversed by Au treatment (Fig. 6E and F). Cleaved caspase 3 fluorescence staining results were consistent with those obtained using western blot (Fig. 6G and H). We then used TBHP to activate PC12 cells *in vitro* to mimic the influence of oxidative stress on neuronal survival after SCI. CCK8 detection and live-dead labeling can both show that Au treatment clearly reversed TBHP-induced neuronal apoptosis (Fig. 7B, G and H). Additionally, after Au therapy, the amount of mitochondrial anti-apoptotic protein (Bcl-2) was partially restored (Fig. 7C-F). These results suggested that Au protected neurons from mitochondrial dysfunction-induced apoptosis.

Previous studies have shown that Au can ameliorate secondary injury by reducing inflammation and neuronal apoptosis caused after SCI. The healing and expansion of neuronal axons is facilitated by successful control of secondary injury [4, 48], and axon regeneration is thought to play a critical role in the recovery of motor function following SCI [9, 61]. Therefore, we evaluated axonal recovery following SCI. Our findings showed that the expression of positive MAP2 axon structural proteins was higher, the region was more densely populated with axons, and axonal organization was more regular in the Au group than in the SCI group (Fig. 8B and C). We used immunofluorescence to stain GAP43 (a neurofilament marker), and showed that Au therapy increased GAP43-positive fibers in the lesion margin of SCI and extended toward the lesion center (Fig. 8D and E). When compared to the SCI group, MAP2 and GAP43 protein levels were partially restored in the Au group (Fig. 8F-H). These results indicated that treatment with Au stimulated regeneration of MAP2-positive axons and GAP43-positive neurofilaments by preventing secondary injury after SCI.

There were several limitations to our study. First, the long-term negative effects of Au injections are unknown. Second, Au is believed to exert a strong anti-inflammatory effect and to promote decreased expression of inflammatory mediators released by inflammatory cells via a variety of mechanisms, but additional mechanisms were not evaluated in this study. Furthermore, because several Au receptors have yet to be identified, it is uncertain whether the anti-inflammatory effects on different phenotypes of inflammatory cells and neuroprotective effects on neurons are mediated by different receptors.

## 5. Conclusion

In conclusion, our data showed that Au can promote recovery from SCI by blocking the TLR4/NF-κB pathway to regulate microglial polarization, thereby reducing inflammation and inhibiting neuronal apoptosis mediated by mitochondrial dysfunction.

## Acknowledgements

This work was sponsored by the “Double Thousand Plan” of Jiangxi Province, the National Natural Science Foundation of China (Grant No. 81860472), Nature Science Foundation of Jiangxi Province (Grant No. 20192ACBL21041; 20202BAB206044).

